# Modeling the Influence of Bandwidth and Envelope on Categorical Loudness Scaling

**DOI:** 10.64898/2026.03.30.715393

**Authors:** Stephen T. Neely, Sara E. Harris, Joshua J. Hajicek, Erik A. Petersen, Yi Shen

## Abstract

In a loudness-matching paradigm, a reduction in the loudness of sounds with bandwidths less than one-half octave compared to a tone of equal sound pressure level has been observed previously for five-tone complexes at 60 dB SPL centered at 1 kHz. Here, this loudness-reduction phenomenon is explored using band-limited noise across wide ranges of frequency and level. Additionally, these measurements are simulated by a model of loudness judgement based on neural ensemble averaging (NEA), which serves as a proxy for central auditory signal processing.

Multi-frequency equal-loudness contours (ELC) were measured for each of the adult participants (N=100) with pure-tone average (PTA) thresholds that ranged from normal to moderate hearing loss using a categorical-loudness-scaling (CLS) paradigm. Presentation level and center frequency of the test stimuli were determined on each trial according to a Bayesian adaptive algorithm, which enabled multi-frequency ELC estimation within about five minutes of testing. Three separate test conditions differed by stimulus type: (1) pure-tone, (2) quarter-octave noise and (3) octave noise. For comparison, loudness judgements for all three stimulus types were also simulated by the NEA model, which comprised a nonlinear, active, time-domain cochlear model with an appended stage of neural spike generation.

Mid-bandwidth loudness reduction was observed to be greatest at moderate stimulus levels and frequencies near 1 kHz. This feature was approximated by the NEA model, which suggests involvement of an early stage of the central auditory system in the formation of loudness judgements.

## Introduction

The loudness of a band-limited stimulus varies with stimulus bandwidth even when its physical intensity remains constant. Such dependence, though reported in previous studies, has not been fully understood. The current study examines this phenomenon across wide ranges of stimulus levels and frequencies and in adults with normal hearing (NH) and hearing loss (HL).

For a band-limited stimulus, as the stimulus bandwidth increases beyond one certain critical bandwidth, the loudness increases. This phenomenon is known as spectral summation of loudness (Leibold et al., 2007). Classic models of loudness (e.g., Moore et al., 1997) interpret the spectral summation in terms of peripheral tuning. That is, the peripheral auditory system can be modeled as a band of bandpass auditory filters, resembling the frequency tuning at various cochlear locations. For a stimulus with a bandwidth entirely within an auditory filter, the excitation at the output of that filter dictates the loudness. When the bandwidth of the stimulus increases such that it excites more than one auditory filter, the loudness is based on the summation of the outputs from these individual filters, leading to an increase in loudness.

Such theories of loudness are challenged by the observation that a pure tone and a narrowband complex stimulus, of equivalent physical intensity and both within a single auditory-filter width, may have a difference in loudness. For example, Zwicker & Fastl (1999), and later replicated by Zhang & Zeng (1997), reported that to match the loudness of a two- or three-tone complex centered at 1 kHz with the overall bandwidth less than 10 Hz, a 1-kHz reference tone would have to be presented about 3 dB higher than the sound pressure level of the complex.

Leibold et al., (2007) reported similar lower loudness for a within-auditory-filter five-tone complex at 1 kHz compared to a 1 kHz pure tone of the same level, with a 5-dB level difference between the two types of stimuli at equal loudness. Hots et al. (2014) systematically examined the effect of bandwidth on loudness using band-limited noises and reported a non-monotonic dependence of loudness on bandwidth. When the bandwidth was within a single auditory filter, the loudness decreased with increasing bandwidth; as the bandwidth extended beyond the auditory-filter width, the loudness increased due to spectral summation. These authors have termed the observed non-monotonicity as “mid-bandwidth loudness depression” (MBLD). Hots et al. (2014) further reported that MBLD was more evident at lower presentation levels. The phenomenon was observed at all three stimulus center frequencies tested (750, 1500, and 3000 Hz), the bandwidth where the greatest loudness reduction was found tended to shift higher for higher center frequencies. Hots et al. (2014) replicated this finding using both loudness matching and categorical loudness scaling (CLS) tasks. Using band-limited noise, Hots et al. (2016) observed MBLD in adult participants with HL. The degree of the loudness reduction was less than in NH adults, potentially due to the high presentation levels required for the participants with HL.

Although the underlying mechanisms for MBLD, especially the loudness reduction with bandwidth for narrowband stimuli within an auditory filter, have not been fully understood, some potential alternative hypotheses have been discussed previously (Hots et al., 2014). These include (1) cochlear compressive nonlinearity and (2) temporal envelope processing.

Compressive nonlinearity may lead to an effective reduction of the total output from the cochlea as the stimulus bandwidth increases. This hypothesis has been initially explored by Hots et al. (2016) using the dual-resonance nonlinear (DRNL) model of the auditory periphery. Although the model was able to produce a suppression effect, it cannot fully explain the observed loudness reduction. It is possible that the subtle response dependence on stimulus bandwidth produced by peripheral nonlinearity may be amplified by certain central mechanisms, leading to a greater degree of effect on loudness than a peripheral model can predict.

Alternatively, it has been hypothesized that the MBLD may be related to how the auditory system processes temporal-envelope information. Band-limited stimuli typically exhibit fluctuations in their temporal envelopes, or amplitude modulation. More importantly, as the stimulus bandwidth decreases, the average modulation rate of the envelope would decrease. Therefore, the dependence of loudness on stimulus bandwidth can also be considered as a dependence on modulation rate. Zhang & Zeng (1997) have suggested that the loudness reduction with increasing stimulus bandwidth within the auditory filter width can be interpreted by the low-pass characteristics for temporal modulation processing, similar to that observed using a modulation detection task (e.g., Viemeister, 1979). More recent loudness models with a central temporal integration stage (e.g., Moore et al., 2016) were able to capture certain aspects of the phenomenon (Rennies et al., 2010), with noticeable discrepancies (Hots et al., 2013).

The present study explores the MBLD phenomenon for band-limited noise across a wide range of levels and frequencies using categorical-loudness-scaling (CLS). Results are interpreted by simulating loudness judgements with a computational model of the auditory system.

Importantly, the computational model combines active, nonlinear cochlear mechanics with the generation of neural spike rates in auditory nerve and a rudimentary central auditory processing stage in the form of neural ensemble averaging (NEA).

## Methods

### Participants

Two groups of adults were recruited from the greater Omaha metropolitan area via a database of volunteers who agreed to be contacted about research that is maintained by Boys Town National Research Hospital (BTNRH). These included a group of 32 adults with NH and a group of 68 adults with HL. The participants in the NH group were between 19 and 69 years of age (mean: 37.7 years; standard deviation: 15.3 years) and had pure-tone average thresholds (PTA, defined as an average hearing threshold across 0.5, 1, and 2 kHz) less than or equal to 15 dB HL. The participants in the HL group were between 32 and 83 years of age (mean: 61.4 years; standard deviation: 12.6 years) and had PTA thresholds between 16 and 55 dB HL. Testing was completed monaurally. For all participants, the ear with lower PTA was selected as the test ear. If there was no difference between ears, then the test ear was selected randomly with an effort to include equal numbers of right and left ears. Across all participants, the test ear PTAs were between 10 and 73 dB HL (mean: 27.9 dB HL; standard deviation: 13.2 dB). It should be noted that the grouping based on PTA (0.5, 1, and 2 kHz) does not account for potential hearing loss at higher frequencies (e.g., 4–8 kHz). While unnecessary for categorization, high-frequency thresholds may influence the loudness perception of wideband stimuli.

Participants provided informed consent before data collection. The experimental protocol was approved by the BTNRH Institutional Review Board.

### Stimuli

In separate conditions, the stimuli were either pure tones, quarter-octave noises, or octave noises. The noises were generated in the frequency domain, with the magnitudes within the passband drawn from a Rayleigh distribution and the phases drawn from the uniform distribution ranging from 0 to 2π. The stimulus magnitudes outside of the passband were zero-padded, before obtaining the time-domain waveform via the inverse Fourier transform. The stimulus duration was 1 s, including 200-ms raised-cosine ramps at the onset and offset. The stimulus level and center frequency were determined before each CLS trial by a Bayesian adaptive procedure, namely the quick-Categorical-Loudness-Scaling or qCLS procedure, as specified in Shen et al. (2025). All stimuli were generated digitally with a sampling rate of 44.1 kHz, presented to the test ear via an audio interface (RME Babyface Pro) and an insert earphone (Etymotic Research ER3A). The qCLS procedure covered a stimulus center frequency range of 0.25 to 6 kHz and a dynamic level range typically spanning 0 to 100 dB SPL, subject to individual safety limits.

### Procedure

For each participant, the experiment was conducted in a single session. The participant sat in a double-walled sound attenuating booth during the experiment. Verbal and written instructions of the CLS task were provided to the participants before the qCLS runs, however no prior training or practice was conducted. Upon presentation of each narrowband stimulus, participants rated its loudness using the 11-category specified in ISO 16832:2006 (International Organization for Standardization, 2006).

A graphical display of 11 loudness categories appeared on a computer monitor as colored horizontal bars scale (see Fig. 1). Seven of the bars had meaningful text labels (“Can’t Hear,” “Very Soft,” “Soft,” “Medium,” “Loud,” “Very Loud,” and “Too Loud”) but participants were encouraged to use all 11 bars. The label ‘Too Loud’ was selected to maintain consistency with historical data collection protocols at this research center, noting that ISO 16832 specifies ‘Extremely Loud’ for this category. After the presentation of each stimulus, the participant selected the category that best represented their perception of the loudness of the sound.

**Figure 1.**
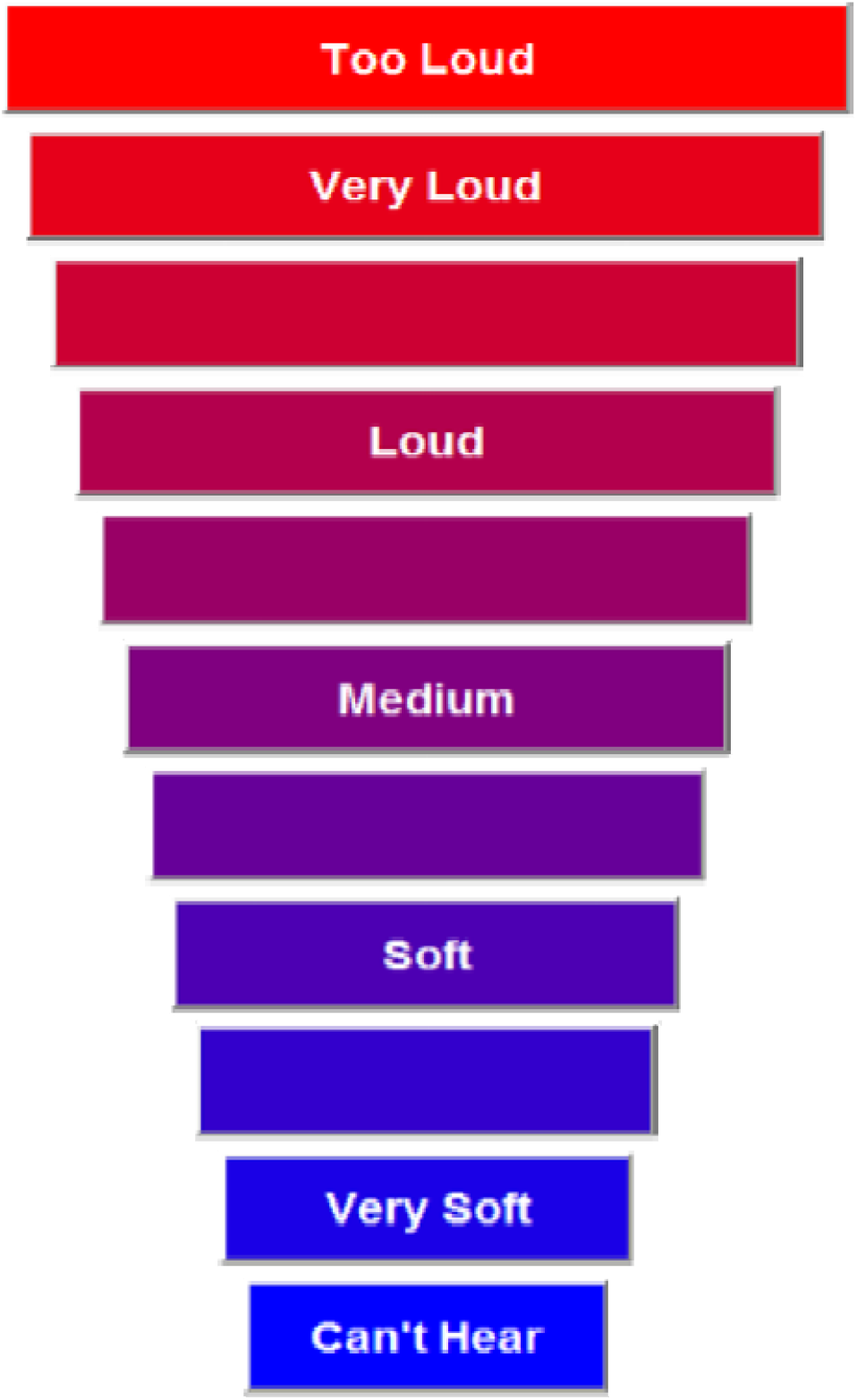
The CLS procedure (ISO16832) assigns meaningful labels to loudness categories that are selected by listeners to describe with various levels, bandwidths, and frequencies. Test efficiency was improved by an adaptive-tracking algorithm (Shen et al. 2024) that applies a Bayesian procedure based on an empirical set of multiple-category psych metric functions.

The three stimulus conditions were tested in a random order. For each condition, a run of the qCLS procedure was conducted. Each qCLS run consisted of 100 category loudness scaling trials and took approximately five minutes to complete. The qCLS procedure determined the stimulus level and center frequency before each trial in an adaptive manner to facilitate sampling stimuli within the participant’s dynamic range. The efficiency of the qCLS method (approximately 5 minutes per run) relies on a Bayesian adaptive procedure that updates a global probability distribution across both frequency and level dimensions simultaneously. Unlike ISO 16832, which treats frequencies independently, qCLS assumes a correlated loudness surface to expedite data collection.

Following data collection, the trial-by-trial stimuli and responses from each qCLS run were used to fit a data-driven model, which determined the loudness profile of the corresponding listener (see Shen et al. 2025 for the computational details). The loudness profiles were reported as the 10 category boundaries separating the 11 response categories across the 10 frequencies (0.25, 0.5, 0.75, 1, 1.5, 2, 3, 4, 6, and 8 kHz) predicted by the fitted data-driven model. Note that although the qCLS procedure did not sample stimuli at 8 kHz due to transducer limitations, the fitted data-driven model nevertheless allowed predicting the category boundaries and equal-loudness contours at 8 kHz.

### Statistical Analysis

To assess the effects of hearing status and stimulus parameters on loudness perception, a linear mixed-effects model (LMM) was fitted to the measured category boundary levels (dB SPL). The model structure included fixed effects for **Hearing-Loss Group** (Normal Hearing vs. Hearing Loss), **Stimulus Bandwidth** (Pure Tone, Quarter Octave, One Octave), and **Loudness Category** (ten inter-category boundaries), as well as their interactions. To account for the repeated-measures design, a random intercept was included for each **Participant**. The model was fitted using the Restricted Maximum Likelihood (REML) method in MATLAB (MathWorks, Natick, MA). Type III F-tests were calculated using Satterthwaite’s approximation for degrees of freedom. Comparisons of loudness growth slopes were performed using independent-samples t-tests.

## Results

### CLS Profiles

The present results combine measurements on 100 participants (32 with normal hearing and 68 with slight, mild or moderate hearing loss) who were assigned to hearing-loss categories based on the PTA of their hearing thresholds. Median loudness profiles of participants with hearing loss are shown in Fig. 2. The median audiometric thresholds of participants with hearing loss are shown as squares in lower-left panel for comparison.

**Figure 2.**
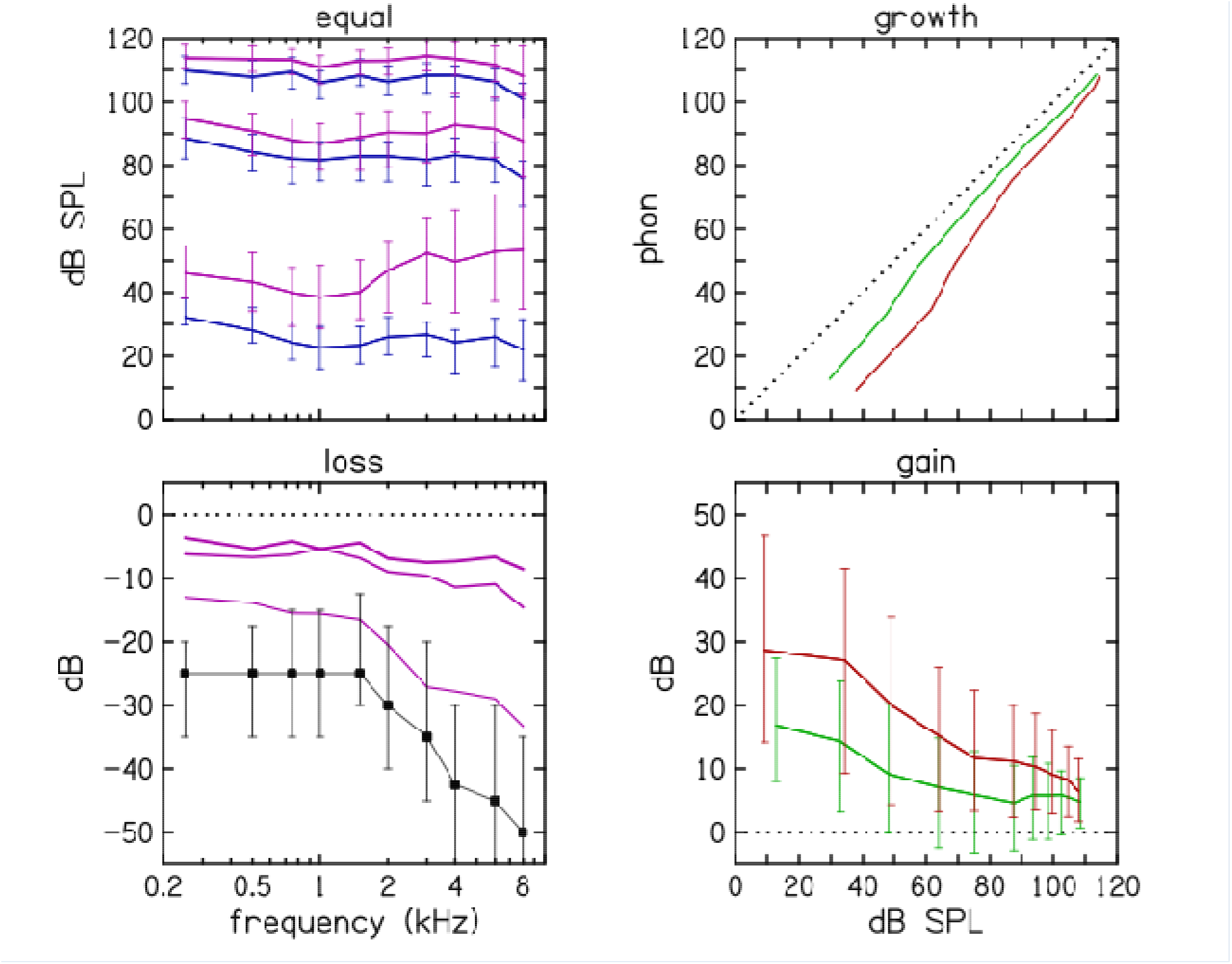
Loudness profile medians across participants with normal hearing and hearing loss. Loudness is plotted in four different ways: (1) Equal loudness contours (ELCs, upper-left panel) for three loudness categories (very soft, medium, very loud) are shown for participants with both normal hearing (blue) and hearing loss (magenta). (2) Loudness loss (lower-left panel, magenta) subtracts the *average* normal-hearing ELCs from the median hearing-loss ELCs. Median hearing thresholds of participants with hearing loss (black squares) are shown for comparison. (3) Loudness growth (upper-right panel) shows median normal-hearing loudness on the vertical axis versus median hearing-loss loudness at 1 kHz (green) and 4 kHz (red). (4) Hearing-aid gain (lower-right panel) replots loudness growth as the gain required to restore normal loudness for each frequency as functions of stimulus level. Error bars represent inter-quartile ranges.

### Loudness Growth and Recruitment

To address the dependence of loudness growth on hearing status across the entire dynamic range, a linear mixed-effects model was fitted to the data. The analysis revealed a significant interaction between **Hearing-Loss Group** and **Loudness Category** (*F*(9,29601) = 51.74, *p* < .001), indicating that the separation between normal-hearing and hearing-impaired functions is not constant but depends on the loudness level. This was further quantified by analyzing the slope of the loudness functions (change in categorical units per dB); the hearing-impaired group demonstrated significantly steeper slopes (*M* = 0.12 cat/dB) compared to the normal-hearing group (*M* = 0.10 cat/dB), *t*(98) = 3.05, *p* = .003). This steeper growth effectively compresses the auditory dynamic range, consistent with the loss of cochlear nonlinearity associated with sensorineural hearing loss. On average, hearing-impaired listeners experience steeper loudness growth below 80 dB SPL for all stimulus bandwidths (Fig 3, left panel).

**Figure 3.**
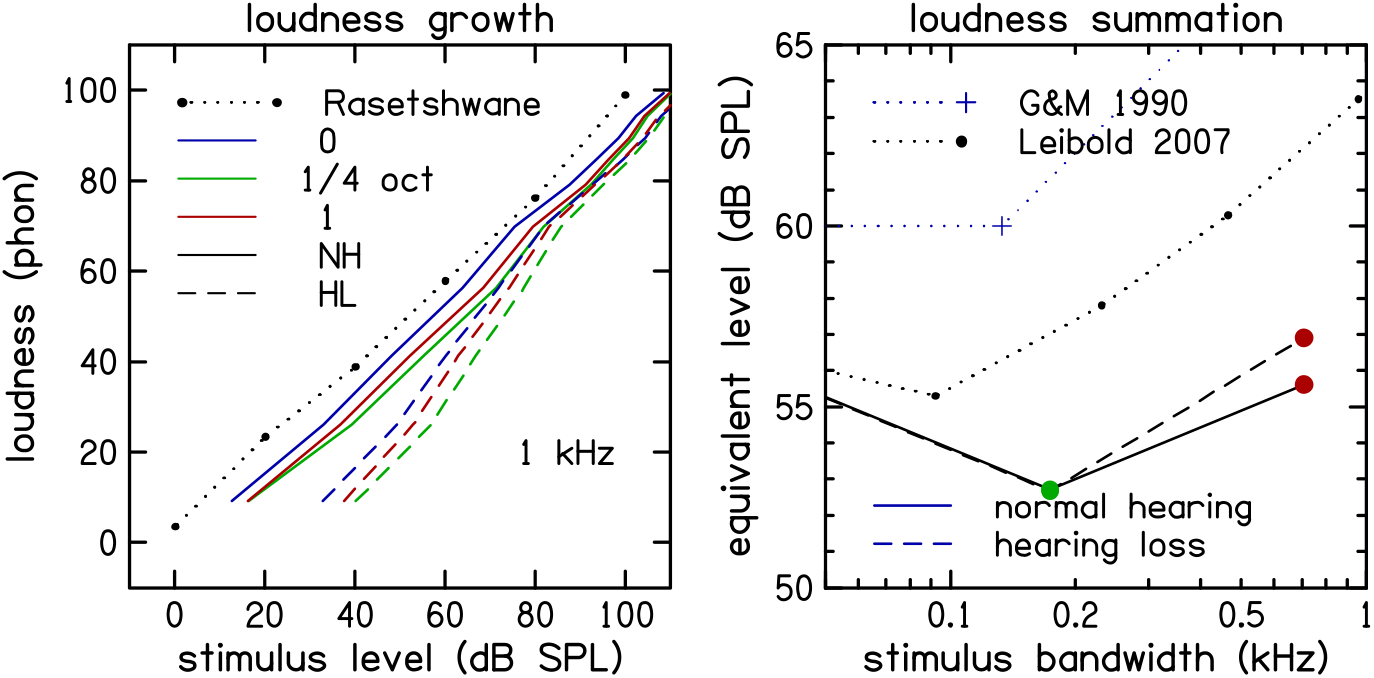
Loudness growth at 1 kHz as a function of stimulus level (left panel) averaged across participants with normal hearing (NH, solid lines) and hearing loss (HL, dashed lines). NH loudness growth of tones (solid blue line). The loudness growth of quarter-octave (green) and one-octave (red) band-limited noise have nearly the same slope. HL loudness is shifted to the right. The sound pressure level (SPL) of an equally loud tone (right panel) is shown for quarter-octave (green) and one-octave (red) stimuli at 60 dB SPL.

### Mid-Bandwidth Loudness Depression

To investigate the MBLD, a focused analysis was conducted at the critical frequency of 1 kHz. This revealed that the equivalent level for quarter-octave noise was significantly higher than for pure tones for both the NH group (*β =* 4.08 dB, *p* < .001) and the HL group (*p* < .001). Consistent with the hypothesis that sensorineural hearing loss compresses the dynamic range, the magnitude of this reduction was significantly larger for NH listeners compared to HL listeners (**Group** × **Bandwidth interaction:** *p* = .042).

Looking across bandwidth at 60 dB SPL (Fig. 3, right panel) reveals that the loudness of quarter-octave noise was about 7 dB less than tones and about 3 dB less than one-octave noise loudness data from Leibold et al. (2007) and loudness prediction from Moore et al. (1997) and are plotted for comparison. Looking across stimulus frequency and level (Fig. 4, left panel), we observe quarter-octave loudness reduction is greatest at 1 kHz and 60 dB SPL but extends over a range of frequencies and levels.

**Figure 4.**
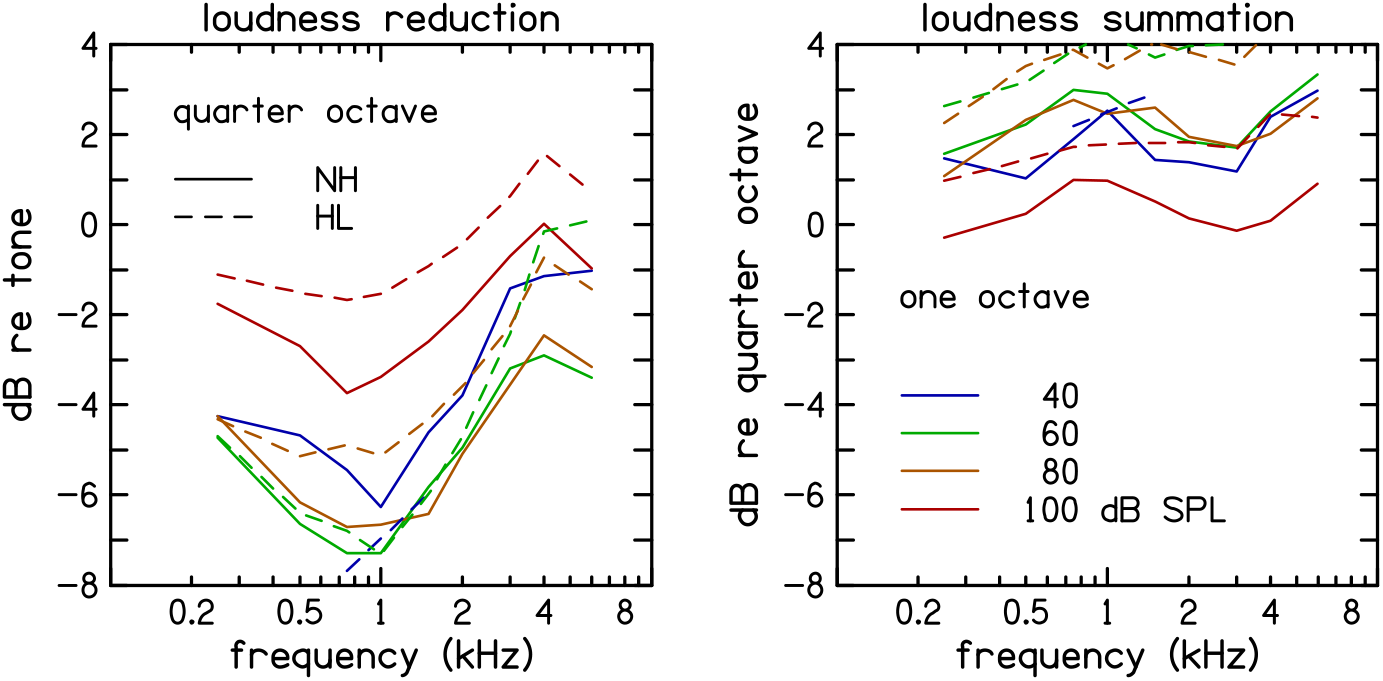
Influence of bandwidth on loudness at four stimulus levels. *Loudness reduction* (left panel) is defined as the change in loudness when quarter-octave noise is compared to a tone at the same stimulus level and center frequency. *Loudness summation* (right panel) is defined as the change in loudness when one-octave noise is compared to a quarter-octave noise at the same stimulus level and center frequency. Stimulus level is represented by different colors. Averages across participants with normal hearing (solid lines) and hearing loss (dashed lines) are superimposed.

### NEA Model

Loudness was modeled by appending neural spike-rate and ensemble-averaging stages to an active, nonlinear, time-domain model of cochlear mechanics (Neely & Rasetshwane, 2017). Neural-spike generation included a diffusion model of synaptic adaptation to mimic forward-masking effects. Neural-ensemble averaging was implemented by sandwiching a partial-loudness summation between spike-rate compression and perceptual expansion:

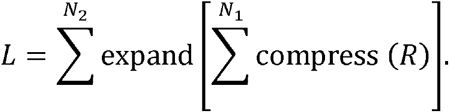

In this equation, R = single-nerve-fiber spike rate, N_1_ = number of nerve fibers per ensemble, N_2_ = number of ensembles in auditory nerve, and L = loudness (sones). Perceptual expansion refers to compensation by the central auditory system that effectively corrects for spike-rate compression in the peripheral auditory system. While computationally intensive, a time-domain model of cochlear mechanics was selected over a functional model to specifically test the hypothesis that fast-acting peripheral compression, coupled with synaptic adaptation, could contribute to MBLD.

Four types of narrow-band stimuli (Fig. 5) that differed in amount of envelope fluctuation were presented as inputs to this loudness model. Single-trial estimates of the modeled loudness for the narrow-band stimuli are shown in Fig. 6 and compared with the loudness measurements from the present study (see Fig. 3). The results show more mid-bandwidth loudness reduction for stimuli with more fluctuation in their temporal envelope. The amount of mid-bandwidth loudness reduction in the model for the “five-tone complex” and “flat noise” stimuli agreed with corresponding experimental observations (Leibold et al., 2007; present study).

**Figure 5.**
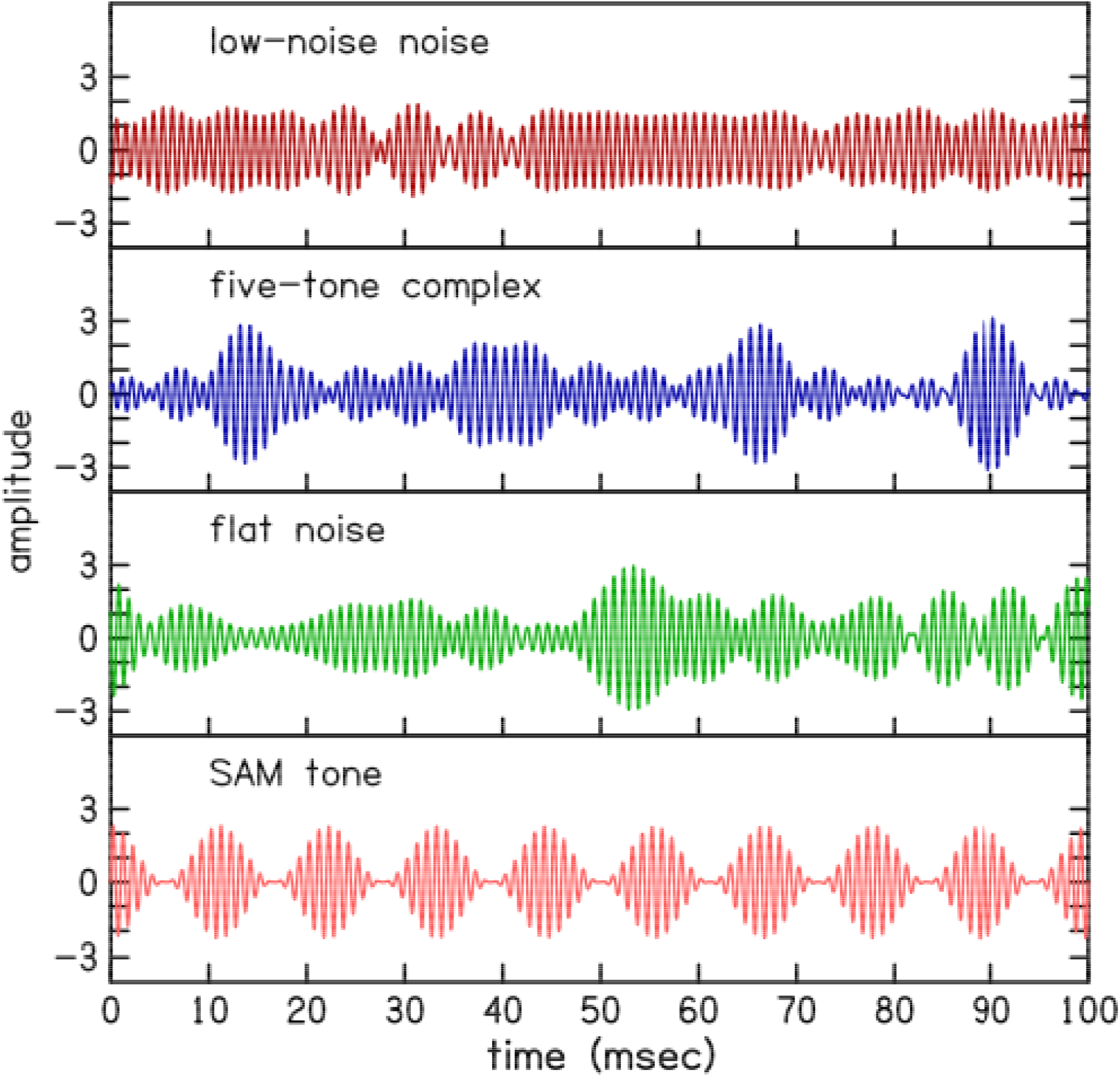
These waveforms represent four different stimulus types that have the same frequency (1 kHz), level, and bandwidth (quarter-octave) but differ in their envelope fluctuation. The rms amplitude of each waveform is equal to 1.

**Figure 6.**
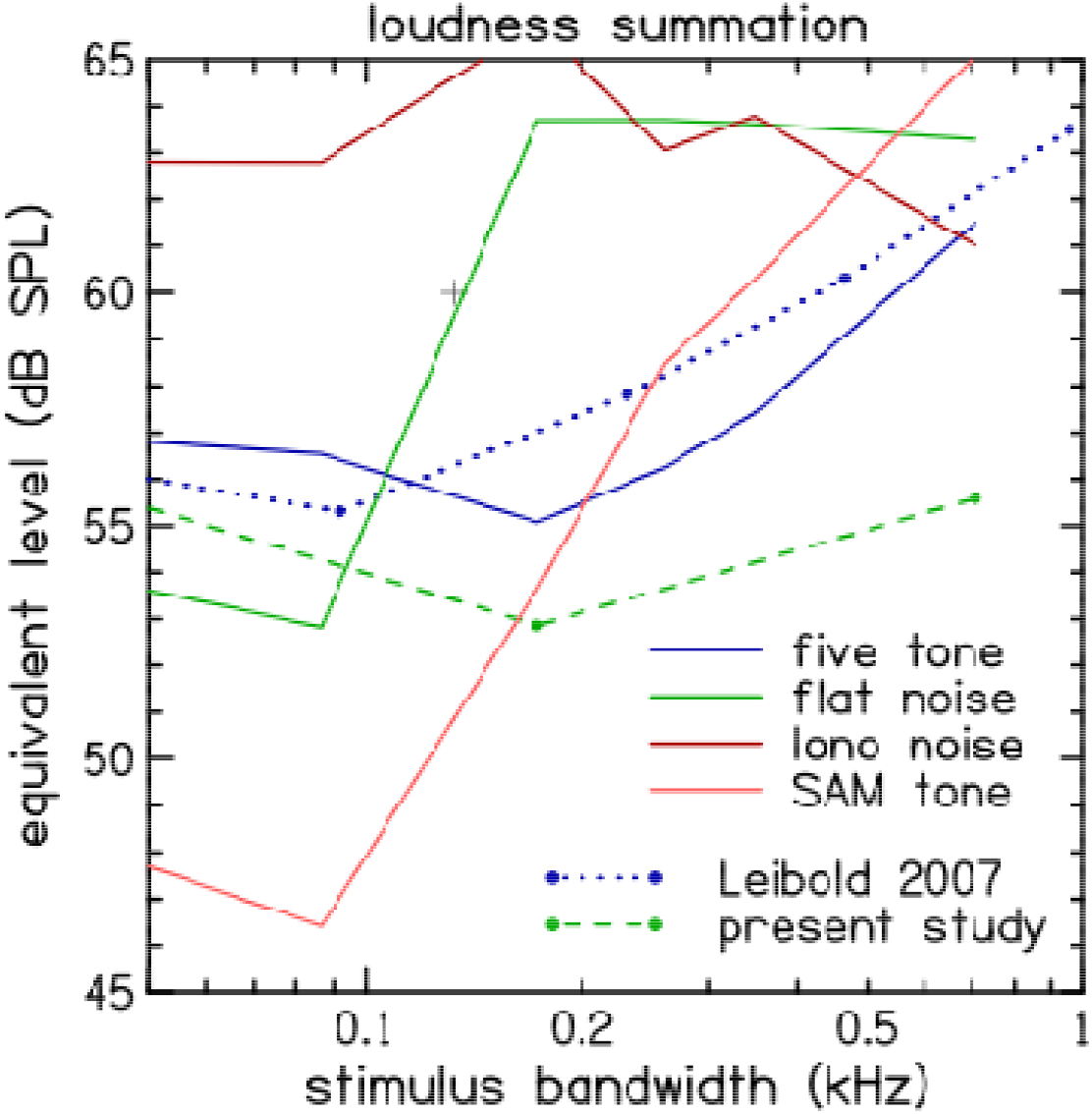
Influence of stimulus type on narrow-band loudness reduction in a nonlinear, active model of cochlear mechanics. The loudness predictions of the model (solid lines) are shown for four different stimulus types: five-tone complex (blue), band-limited noise (green), low-noise noise (magenta), and SAM tone (orange). For comparison, the loudness measurements in the present study used only bandlimited noise stimuli (green, dashed line). Whereas loudness measurements in the Leibold et al. (2007) study (blue, dotted line) used only five-tone complex stimuli.

## Discussion

The quarter-octave loudness reduction observed in this study in CLS measurements has previously been called MBLD and observed for narrow-band-noise stimuli in both normal-hearing and hearing-impaired listeners (Hots et al., 2014; 2016) in both loudness matching and scaling procedures.

The magnitude of the MBLD observed in the present study is generally consistent with, though slightly larger than, the magnitude reported in previous literature. We observed a maximum loudness reduction of approximately 7 dB for quarter-octave noise compared to pure tones at 1 kHz. This aligns with, but exceeds, the approximately 5-dB effect reported by Leibold et al. (2007) using a loudness-matching paradigm with five-tone complexes. Regarding level dependence, our results indicated that MBLD was most pronounced at moderate presentation levels (60 dB SPL). This contrasts somewhat with Hots et al. (2014), who reported that MBLD for band-limited noise was more evident at lower presentation levels. This discrepancy may reflect differences in the measurement tasks (adaptive qCLS versus adaptive categorical scaling or matching) or the spectral characteristics of the noise stimuli used. Additionally, consistent with Hots et al. (2016), we observed that hearing-impaired listeners generally exhibited a reduced MBLD effect compared to normal-hearing listeners, likely due to the recruitment associated with sensorineural hearing loss which compresses the dynamic range and steepens loudness growth functions.

The statistical confirmation of steeper loudness slopes in the HL group (*p* = .003) quantitatively supports the existence of recruitment. While the statistical analysis confirmed that MBLD remains significant in hearing-impaired listeners (*p* < .001), the significant Group × Bandwidth interaction (*p* = .042) demonstrates that the magnitude of this loudness reduction is effectively compressed by the hearing loss.

Loudness reduction in the model was a consequence of (1) fast-acting compression in the signal delivered to the auditory nerve and (2) neural-ensemble averaging in the formation of partial loudness judgements, and (3) perceptual expansion of prior to final loudness judgements. Neural ensembles are subsets of auditory nerve fibers that originate from a region of the cochlear partition that spans about 2 mm. For comparison, a critical band spans about 0.9 mm. Averaging across such ensembles, which presumably occurs within the cochlear nucleus, provides a fast-acting averaging mechanism for noise reduction that complements averaging of neural signals across time.

Our pilot modeling efforts have suggested that the amount of loudness reduction in the model is sensitive to the parameter that specifies the ensemble width. This observation supports the view that ensemble averaging may be an essential feature of auditory signal processing. Likewise, the robust observation of mid-bandwidth loudness reduction is consistent with the mechanism of neural-ensemble averaging as predicted by the model. However, because the high computational cost of solving active, nonlinear cochlear mechanics in the time domain, combined with the stochastic requirements of generating stable post-stimulus time histograms for neural ensembles, hindered comprehensive exploration of alternative parameter values, the present model results should not be considered the best that could be achieved by this approach. For example, model computation times may extend several days even for a subset of desired stimulus conditions and shortened stimulus durations. Future modeling efforts will use models that require less computation time. Future research could systematically test neural ensemble averaging as an underlying mechanism for MBLD and to examine additional alternative explanations.

The observation of MBLD in so many participants across large ranges of stimulus frequencies and levels would not have been feasible without the efficiency of the CLS procedure combined with our multi-frequency, adaptive tracking, qCLS method (Shen et al., 2025).

## Conclusions

Quarter-octave loudness reduction is thought to be due to a combination of factors, including stimulus *envelope fluctuation, fast-acting compression* in cochlear mechanics, and *neural-ensemble averaging*. A signal-processing model that replicates key features of loudness reduction implicates temporal-envelope fluctuation as being the signal property responsible for this phenomenon. The demonstrated feasibility of the current model to qualitatively predict MBLD points to the importance of considering both peripheral and central mechanisms when modeling loudness perception.

## Acknowledgement

We are very thankful to the patients and the clinical teams of Radboud University Nijmegen Medical Centre (Netherlands) and Centre Hospitalier Universitaire d’Angers (France) for their participation in this operationally challenging study.

## Acknowledgements

The authors would like to thank Daniel Rasetshwane for his contributions to the development of the CLS methods used in this study.

## Statements and Declarations

The authors declare that they have no significant financial conflicts of interest to disclose.

## Ethical Considerations

This research has been approved by the Institutional Review Board at Boys Town National Research Hospital. All participants provided written informed consent before data collection.

## Data availability statement

The data that support the findings of this study are available from the corresponding author upon reasonable request.

